# An interpretable ML model to characterize patient-specific HLA-I antigen presentation

**DOI:** 10.1101/2023.03.12.532264

**Authors:** Shaoheng Liang, Xianli Jiang, Yulun Chiu, Haodong Xu, Kun Hee Kim, Gregory Lizee, Ken Chen

## Abstract

Personalized immunotherapy holds the promise of revolutionizing cancer prevention and treatment. However, selecting HLA-bound peptide targets that are specific to patient tumors has been challenging due to a lack of patient-specific antigen presentation models. Here, we present epiNB, a white-box, positive-example-only, semi-supervised method based on Naïve Bayes formulation, with information content-based feature selection, to achieve accurate modeling using Mass Spectrometry data eluted from mono-allelic cell lines and patient-derived cell lines. In addition to achieving state-of-the-art accuracy, epiNB yields novel insights into the structural properties, such as interactions of peptide positions, that appear important for modeling personalized, tumor-specific antigen presentation. epiNB uses substantially less parameters than neural networks, does not require hyperparameter tweaking and can efficiently train and run on our web portal (https://epinbweb.streamlit.app/) or a regular PC/laptop, making it easily applicable in translational settings.

## Introduction

After decades of development, immunotherapy is now at the frontier of cancer therapy and has been demonstrated to be beneficial to clinical outcomes in many cancer types [1–8]. The capacity to redirect T-cells against tumors has raised a large degree of interest in identifying patientspecific peptides that can be targeted therapeutically. Human leukocyte antigen (HLA) molecules are cell surface proteins that present peptides derived from intracellular proteins to T cells, triggering immune recognition and activation. HLA Class I (HLA-I) molecules present peptides to CD8+ cytotoxic T cells, which can redirect their killing activity towards tumor cells through recognition of specific tumor-associated HLA-I/peptide complexes. HLA-I molecules typically bind and present peptides with a length range of 8 to 13 amino acids, and collectively display thousands of peptides (the “immunopeptidome”) that represent a snapshot of the current translated cellular proteome (i.e., the collection of all proteins in the cell). HLA binder prediction is important in broad areas, such as cancer vaccine designs [1], adopted cell therapy [2], viral infection [3], autoimmune diseases [4], and organ transplantation [5]. Identifying relevant tumorspecific peptide targets in individual cancer patients is a challenging problem, due to significant genetic, epigenetic and immunological heterogeneity across individual patients, and across celllular populations within each patient.

Advances in next-generation sequencing now allow for typing of personalized HLA haplotypes and for detection of somatic mutations, thus enabling prediction of personalized tumor-specific antigens (TSAs) for use in personalized immunotherapy. Despite the application of large neural networks such as NetMHCpan, predicting TSAs from genomic mutations remain challenging for cancer patients carrying rare HLA alleles, or having altered antigen processing and presentation processes. In comparison, Mass spectrometry (MS)-based profiling of tumor-derived, eluted peptides allows directly observation of personalized immunopeptidome [6]. However, the MS-based assays usually yield only hundreds to thousands of peptides, a small subset of presented peptides, making it difficult to directly detect TSAs, or train machine-learning-based models. Thus, the field urgently calls for new methods that can make accurate predictions based on small size data.

The classical immunopeptidome consists of peptides collectively presented by up to six alleles per patient (i.e., two alleles each of HLA -A, -B, and -C). Peptide-HLA complexes from cells are experimentally captured by monoclonal antibodies (mAbs) to be sequenced. Because allele-specific mAbs are relativley rare, most immunopeptidome studies utilize the pan-HLA-ABC-specific mAb W6/32, which will immunoprecipitate all HLA-I/peptide complexes and thus provide a mixture of peptides eluted from all 6 alleles. To train allele-specific model, the peptide pool must be accurately deconvolved [7], which in itself a challenge when the correspondence of peptides and the alleles are not precisely known [8]. A recent study analyzed peptides eluted from monoallelic cell lines expressing 95 individual HLA-I alleles, providing an invaluable dataset of pure, allele-specific peptides, leading to significantly improved training and performance of HLA-I/peptide binding prediction algorithms [6].

Cancer cells can evade immune surveillance by interfering with the antigen processing and presentation machinery [9]. Known examples include copy number loss of NLRC5, a gene to activate the expression of several components of the antigen presentation process, and overexpression of HSP90, which prevent certain proteins from being processed. There defects are prior to the pHLA binding event and cannot be easily identified using conventional approaches. Hence, training patient specific models on eluded peptides is a more effective way to identify druggable neoantigens.

In addition, the problem is characterized by a lack of negative examples [10], as presented peptides are relatively rare (<1%) and experimental assays often report just binders. Because most machine learning models require negative examples (i.e., non-binders) to train, studies predicting neoantigens generate random, non-specific peptides (either fully random, or stripped from wildtype protein sequences) as negative examples. In addtion, datasets obtained from real applications are extremely imbalanced (<1% positives), leading to a stress test for machine learning techniques.

For antigen presentation, two major classes of methods are widely explored. Allele-specific methods train models on peptides obtained from binding assay or mass spectrometry, while pan-HLA methods aim to find links between HLA sequences and binding peptides and extrapolate them to understudied HLAs. The former approach frames the problem as a classic classification problem, with many kinds of methods being explored, such as expert-system-style scoring systems, decision trees, and eventually an ensemble of them [11]. Recent studies have shown that neural networks (NetMHC-4.0 [12] and HLAthena [6]) generally perform well. The pan-allele methods employ larger models to “translate” between HLA sequences and binding peptides [13–15]. Existing solutions using deep neural networks require large training set and special hardware (GPU), making them less accessible [11,15]. In addition, patient privacy regulations can further restrict uploading of patient data through the web portals of some methods. Moreover, these methods tend to model diversity in the binding step but do not account for endogenous processing and presentation steps (with MHCFlurry [13] being an exception), which make them insufficient for personalized applications.

In this study, we developed epiNB, a Naïve-Bayes-based, semi-supervised, positive-example-only classification method that predicts personalized TSAs from eluded peptide libraries. For feature selection, epiNB employs an unsupervised, mutual-information-based approach and a weakly supervised approach, benchmarked against a Siamese network (**Figure 1**) [16]. It uses a semisupervised pseudo-labeling approach to deconvolve eluted peptides to HLA alleles. Our study revealed the importance of having both single and pairwise amino acid (AA) features in achieving the specificity. As control, we compared epiNB against a variety of state-of-the-art methods of unique breakthroughs in data collection and/or modeling on single-allelic and patient-derived data (**Table 1**; Briefly summarized in **Results**). We show that epiNB achieves comparable or favorable performance, despite of using simpler models. EpiNB is available as a Python package (https://github.com/KChen-lab/epiNB) and an online portal (https://epinbweb.streamlit.app/; **Supplementary Figure 1**).

**Figure 1.**
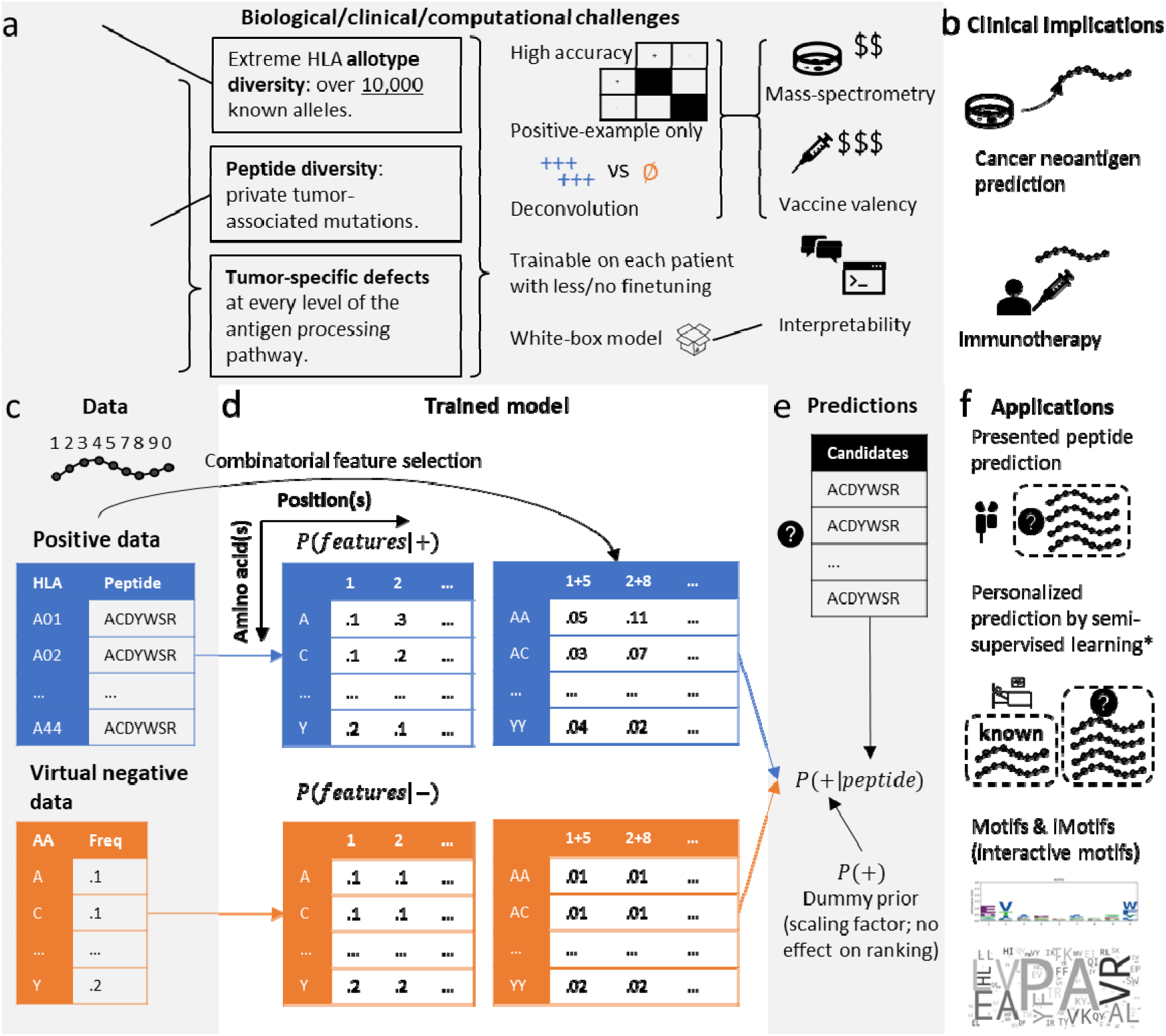
Overview of the epiNB model. (a) Biological, clinical, and computational challenges in predicting personalized immunotherapy target peptides. (b) Clinical applications of MHC-I presented peptide prediction methods. (c-f) Workflow of EpiNB algorithm, which (c) takes positive data (known presented peptides) and prior prior distribution of AAs (equivalent to negative data randomly generated from the distribution) as input and (d) use the frequencies of AAs at each position, as well as combination of positions selected using their mutual information, to (e) make predictions for candidate neoantigens. The method can be used to rank candidate neoantigens, make multi-class classification of HLA origin of a peptide (i.e., deconvolve eluted peptides from patient), and thus retrained on patient specific data to make personalized predictions. As a white-box model, the basis of choosing a peptide, i.e., motifs and iMotifs (interactive motifs) is given, so that clinicians and researchers make more informed judgements.

**Table 1:** Compared methods

| Method | Model | Data | interpretability |
| --- | --- | --- | --- |
| epiNB | Specific | MS | motifs and interactive motifs |
| TransMut [15] (2022) | Pan | MS+BA | Self-attention scores (from transformer) |
| Anthem [11] (2021) | Specific | MS+BA | Black box (ensemble) |
| MHCFlurry [13] (2020) | Pan | MS+BA with deconvolution of processing and binding | Black box (Two NNs for processing and binding) |
| HLAthena [6](2019) | Specific | MS | Black-box (NN) |
| MixMHCpred [7](2022) | Specific | MS with deconvolution of HLA alleles | 1<sup>st</sup> order motifs (from position weight matrices) |
| NetMHC-4.0 [12] (2016) | Specific | BA | Black box (NN) |

## Results

### A personalized semi-supervised positive-example-only peptide classification model

The core of epiNB is a Naïve Bayes classifier modified for positive-example-only learning (**Figure 1c-e**, Methods). In brief, epiNB takes a set of known binders to perform feature selection on the set using mutual information of a combination of peptide positions (**Supplementary Figure 2a**). The binders then serve as the training set to train the model to derive *P*(*feature* | −) for both single positions (referred to as “motifs”)] and interactions of positions (referred to interactive motifs “iMotifs”). The combinations reflect the intrinsic dependency between two peptide residues within an HLA-specific peptide population. For most alleles, the anchor residues (P2 and P0; we number the first 5 amino acids 1, 2, 3, 4, 5, and the last four 7, 8, 9, 0; 0 is also commonly noted as Ω) exhibit low dependency on other residues, indicating their dominant roles in defining the HLA binding (**Supplementary Figure 2a**). Interestingly, the signal of combination of P2 and P0 is the strongest in pan-allele, supporting the importance of having these two positions in determining the HLA binding specificity across alleles. Most machine learning methods require both positive and negative examples to learn, and the common practice in this field is to draw random samples from the prior distribution or human peptidome. However, the Naïve Bayes formulation allows epiNB to directly use the prior distribution of amino acids (AAs) as *P*(*feature*|−) and avoid variances induced by sampling. Using the Bayesian rule, epiNB can then calculate *P*(+|*peptide*). EpiNB includes no hyperparameters to tweak, except for a dummy prior *P*(+), the unconditional probability of observing a binder, which, however, only serves as a scaling factor and has no effect on the ordering of candidate peptides (**Methods**). The model is trained on peptides eluted from cell lines that only have one HLA allele (cf. six alleles of HLA-A/B/C[6], which comprehensively assess the whole antigen presentation process, compared with binding panels, which only consider the binding step (**Figure 1a**).

The result of epiNB can be used to rank potential binders, and to classify the HLA allele a peptide belongs to (**Figure 1f**), both useful in personalized binding peptide prediction, where a few eluted peptides are made available from the patient but can correspond to any of the six alleles of the patient. Blindly training a machine learning model on such a problem is usually suboptimal, because the binders of the six alleles can be vastly different, causing problems for models to draw a reliable classification boundary. To this end, we first use the classification utility of epiNB to assign the peptides to alleles and add the peptides to the training data to refine the model. The final model can be used to make predictions.

Although expert systems have lost ground to machine learning techniques given the booming amount of data, oncologists and immunologists still hold precious knowledge about peptide binding. As an open-box model, epiNB can readily tell the motifs it relies on in deriving the results, which allows practitioners to inspect the rules and make informed decisions.

### EpiNB accurately predicts presented peptides of HLA alleles

To benchmark the performance of epiNB and other methods, we constructed a large-scale test using The Immune Epitope Database (IEDB) (**Figure 2a**) with a total of 1.7M positive peptides across 95 HLA alleles. The experimentally verified binders are used as positive examples (binders) and negative examples (decoys) are generated from the human proteome. Considering that finding binders from candidate peptides are essentially a needle-in-a-haystack problem, we set the ratio of positive and negative examples to 1:99. Following the convention of similar studies [6,13,15], we used both precisions at 40% recall and AUROC as the metric. However, we strongly recommend the precision at 40%, as it is more indicative of the actual performance in vaccine design (**Methods** and **Supplementary Note 1**), and is widely used in recent studies [6,13].

**Figure 2.**
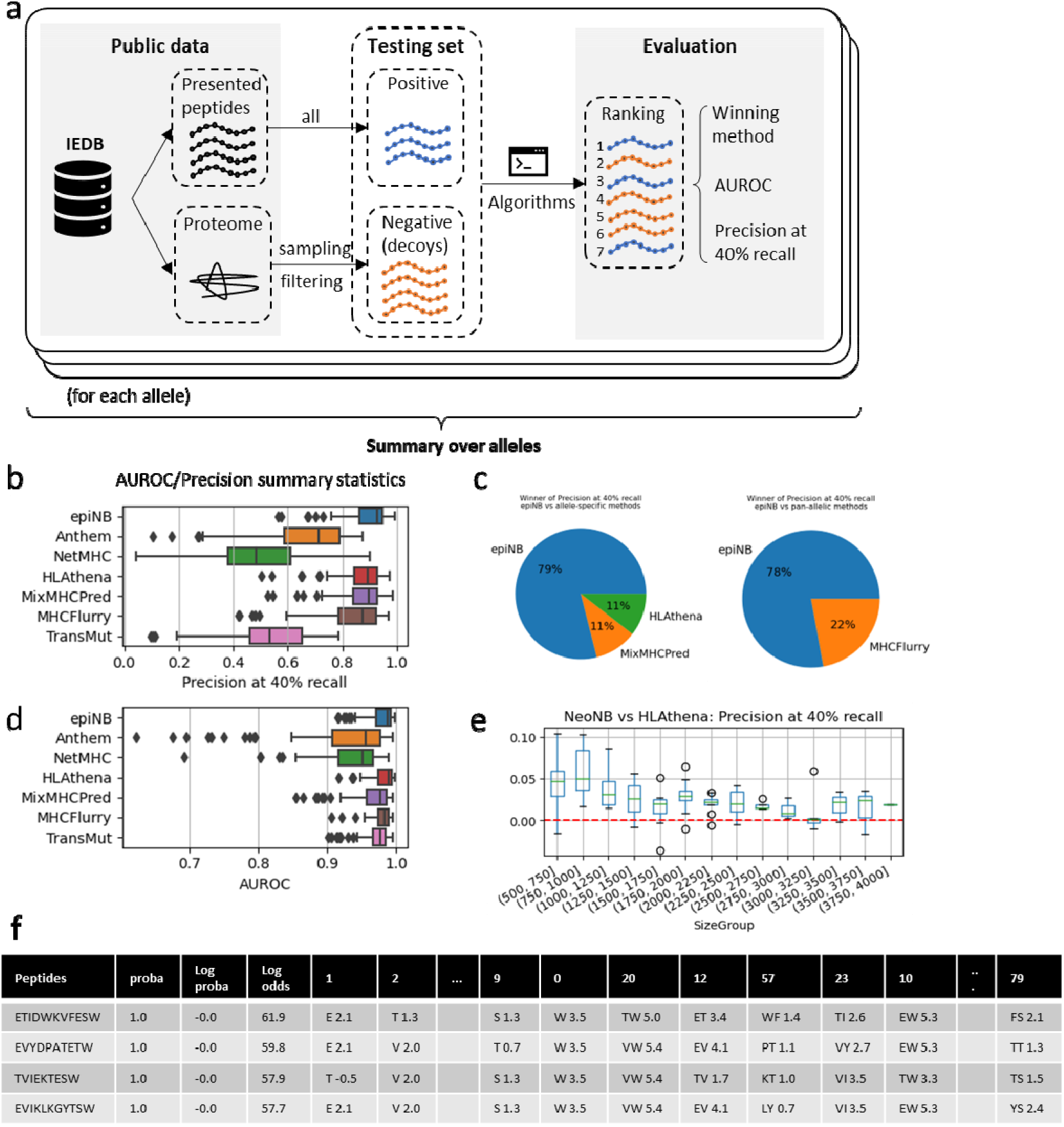
Benchmarking on IEDB dataset. (a) Workflow of benchmarking. (b-c) precision at 40% recall (b) and area under receiver operating curve (AUROC) (c). Higher is better. (d) The proportion of winners over all 95 alleles. More is better. (e) The gain of precision at 40% recall over HLAthena, a neural network method trained on the same dataset. Higher is better. (f) Sample prediction report for A2501. Proba: probability, log proba: log binding probability, log odds: log likelihood of binding minus that of non-binding. The following part of the table shows the AA combinations, and the log odds for it.

For comparison (**Table 1**), we included a variety of state-of-the-art methods of unique breakthroughs in data collection and/or modeling. Four allele-specific models were included. Anthem [11] explored ensemble to optimize the performance of existing data. HLAthena [6] piloted generating a clean training dataset with cell lines with a single HLA allele and used a neural network model for prediction. MixMHCPred [7] automatically deconvolve unlabeled peptidome to generate a large training dataset with position weight matrices for prediction. NetMHC-4.0 is a classic tool that serves as a baseline. We also included two state-of-the-art pan-allelic models. TransMut [15] explored the application of the transformer model in peptide binding prediction and provided self-attention scores as a way to interpret the results. MHCFlurry [13] deconvolved the process of antigen processing and peptide binding to more accurately model the process of antigen presentation. In addition to the (1) experimentally verified binders in IEDB, in the following sections we also compared the methods on (2) eluted peptides from patient-derived cell lines/mouse models, and experimentally verified (3) NSCLC neoantigen and (4) HPV neoantigens, we illustrate that in addition to achieving state-of-the-art accuracy on binder prediction, epiNB also deconvolves peptides from multiple HLA allotypes without manual tuning, leading to accurate modeling of patient data for clinical applications.

EpiNB achieved better precision at 40% recall than other methods (**Figure 2b** and **Supplementary Figure 3a**) on the IEDB datasets with 100 to 10,000 known positive peptides per each of the 95 allele. On nearly 80% of the alleles, epiNB performed better than all other methods (**Figure 2c**). The AUROC is less discriminatory, with epiNB showing a slightly lower score than HLAthena (**Figure 2d** and **Supplementary Figure 3b**). However, individual ROC and Precision-recall plots indicate that epiNB is better in practical scenarios (**Supplementary Figure 4** and **Supplementary Note 1**). To investigate the effect of training data size, we compared the performance difference between epiNB and HLAthena (which are trained on the same dataset) over different training data sizes (**Figure 2e**). EpiNB shows favorable performance across all sizes of training data and especially more gain with smaller training sets.

To fully utilize the interpretability/transparency of the model, we provide various utilities to show binding insights. EpiNB can generate a “prediction report” with the log odds for each position and combinations (**Figure 2f**) to explain the ground of its classification. The entries with high odds (positive and large absolute values) are positive evidence for binding, and the low odds (negative values) are negative evidence. These insights can then be paired with prior knowledge of clinicians and the motif knowledge acquired from inspecting the training process (**Supplementary Figure 5**; more in next subsection), to help clinicians make informed decisions.

### Combinatorial feature selection casts insights into peptide binding

Being one of the most direct ways to inspect the peptides, (1^st^ order) binding motifs of binders have long been studied and well-characterized [6]. Even with predictions made by black-box models, oncologists and immunologists prefer double-checking the motifs. Here, we show that motifs are not always sufficient and focus on the iMotifs found by epiNB, which are nearly as intuitive as motifs but include richer information.

As an example, we show that iMotif with the largest mutual information, P20 (i.e., P2 and P0), in Figure 3. (**Supplementary Figure 5b** shows other pan-allelic combinations and **Supplementary Figure 5c shows** an example of allele-specific combinations.) A high mutual information content indicates that the distribution of the motif would be drastically different from the theoretical distribution assuming independence of the two positions. Thus, we used word cloud to illustrate the abundance of pairs of AAs in the actual and theoretical distribution (**Figure 3a,b**; see **Supplementary Figure 6a,b** for heatmaps and tables conveying the same information), and show the difference between the two distributions in Figure 3c, where PA, VR, LV, and EA show large surplus than they would have been, and VL, TL, and ER show large deficiency. We then check if these combinations are indeed indicative of the alleles. Indeed, we find that about half of peptides from B5601, B5502, B5501, and B5401 feature the PA motif (**Figure 3e**). B5601, B5502, B5501, and B5401 share nearly identical sequences in peptide binding pockets B and F (**Supplementary Figure 6c**; Sequence alignment for B: 9,24,45,63,66,67; F: 74,77,95,97,114,116, 123). The Tyr9, Ile66, and Tyr67 in the binding pocket will preferentially select Pro as the P2 anchor residue to fit its hydrophobic side chain to the hydrophobic cleft formed by Tyr67 and Ile66. The F pocket in these alleles is considered as neutrally charged as only 74 is Asp, therefore showing no selection on Arg or Lys [17], The VR motif, though less prominent than PA, is also highly specific to a group of alleles, such as A3401, where the F pocket is highly negative contributed by Asp74, Asp77, Asp116, and highly selective for Arginine. (**Figure 3f**).

**Figure 3.**
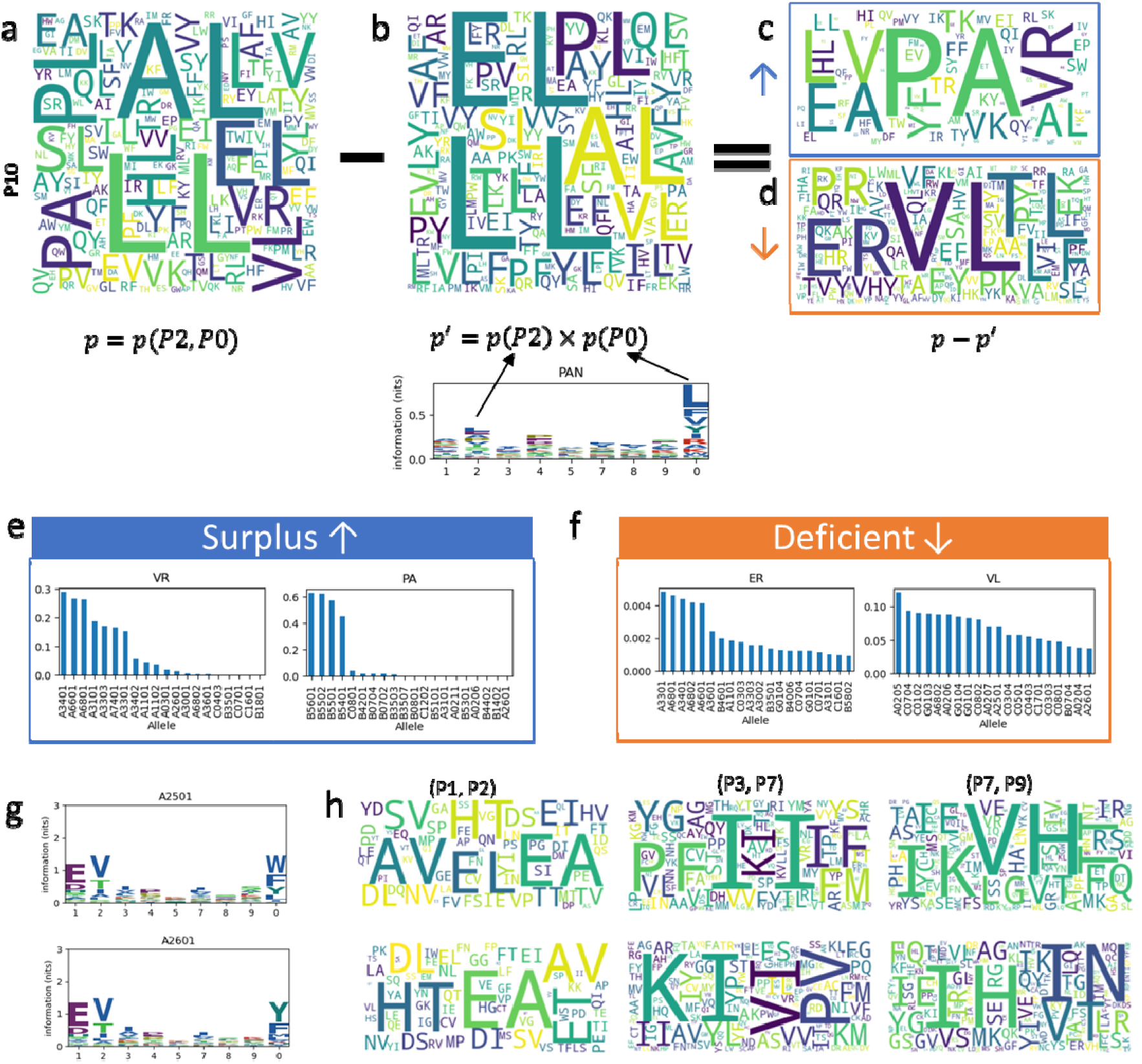
EpiNB improves interpretability of peptide binding predictions. a. Word cloud of observed frequency of iMotifs of the 2^nd^ and last position of the peptides. Larger font size correspond to more occurrences. b. Word cloud of theoretical frequencies of pairs of iMotifs of the 2^nd^ and last position of the peptides, assuming no interaction between the two positions. The logo plot underneath shows the marginal probability of AAs at each position. c-d. The iMotifs that occur more (c, “surplus”) or less (d, “deficient”) frequently than it would have been assuming no interaction between positions. e-f. The top two surplus (e) and deficient (f) iMotifs and the observed frequency of them showing up in the eluted peptides from each allele (only top 20 are shown). g. Logo plots for A2501 and A2601, where the total height of a position is log(20) - entropy(position), and the height of each individual amino acid is proportion to its frequency the position. h. Word clouds for A2501 and A2601 at selected iMotifs.

These insights are an integral part of epiNB, which can be easily retrieved after training. To further illustrate their value, we show the 1^st^ order motifs (**Figure 3g**) and selected combinations (**figure 3h**) for HLA-A2501 and HLA-A2601, which are considered to be in a superfamily [6]. Indeed, the logo plots show very few differences between these two alleles. However, the pan-allelic feature P12 started to reveal some differences, such as “SV” for A2501 and “ET” for A2601, and the allele-specific features P37 and P79 carry even more distinct patterns. These combinations may have similar roots in the protein structure of the HLA complex as P20. It further illustrates that iMotifs carry important information about peptide binding that can be used to interpret the predictions.

We benchmarked the mutual information against a weakly-supervised approach, Siamese network [16], to investigate interactions between peptide positions (**Supplementary Figure 7a**). Siamese network is a deep learning approach to create a low dimensional embedding (**Methods** and **Supplementary Figure 7c**) that is especially suitable for applications on complex inputs that are not suitable for traditional dimensional reduction approaches like PCA. For interpretability, we explicitly included in the network the combinatorial features and record their weights (importance) in generating the embedding (**Supplementary Figure 7b,d**). The result is consistent with the mutual information based approach, reaffirming that the chosen combinatorial features are important characteristics of the peptides. The embedding can also be used to deconvolving patient-derived peptides to specific HLA alleles.

### EpiNB achieves state-of-the-art performance on patient-derived data

We then benchmarked all methods on six datasets derived from cancer patient samples including four pancreatic xenograft samples and two lung cancer surgery samples, each having 300 to 600 known presented peptides. Data in each sample were split into two equal sized training and testing sets, and the testing sets was further amended by 99x more negative examples (**Figure 4a**). For patient data, we took a pseudo-labeling semi-supervised learning approach. Specifically, for each patient, we deconvolved eluted peptides (that may be from any one of six given HLA alleles) and assign them to a specific HLA to augment training data and retrain the model (**Supplementary Figure 8a**). The validity of the deconvolution was verified by the performance of allele classification (**Supplementary Figure 8b**). This approach created patient specific models that integrates the cancer specific defects information underlying the eluted peptides.

**Figure 4.**
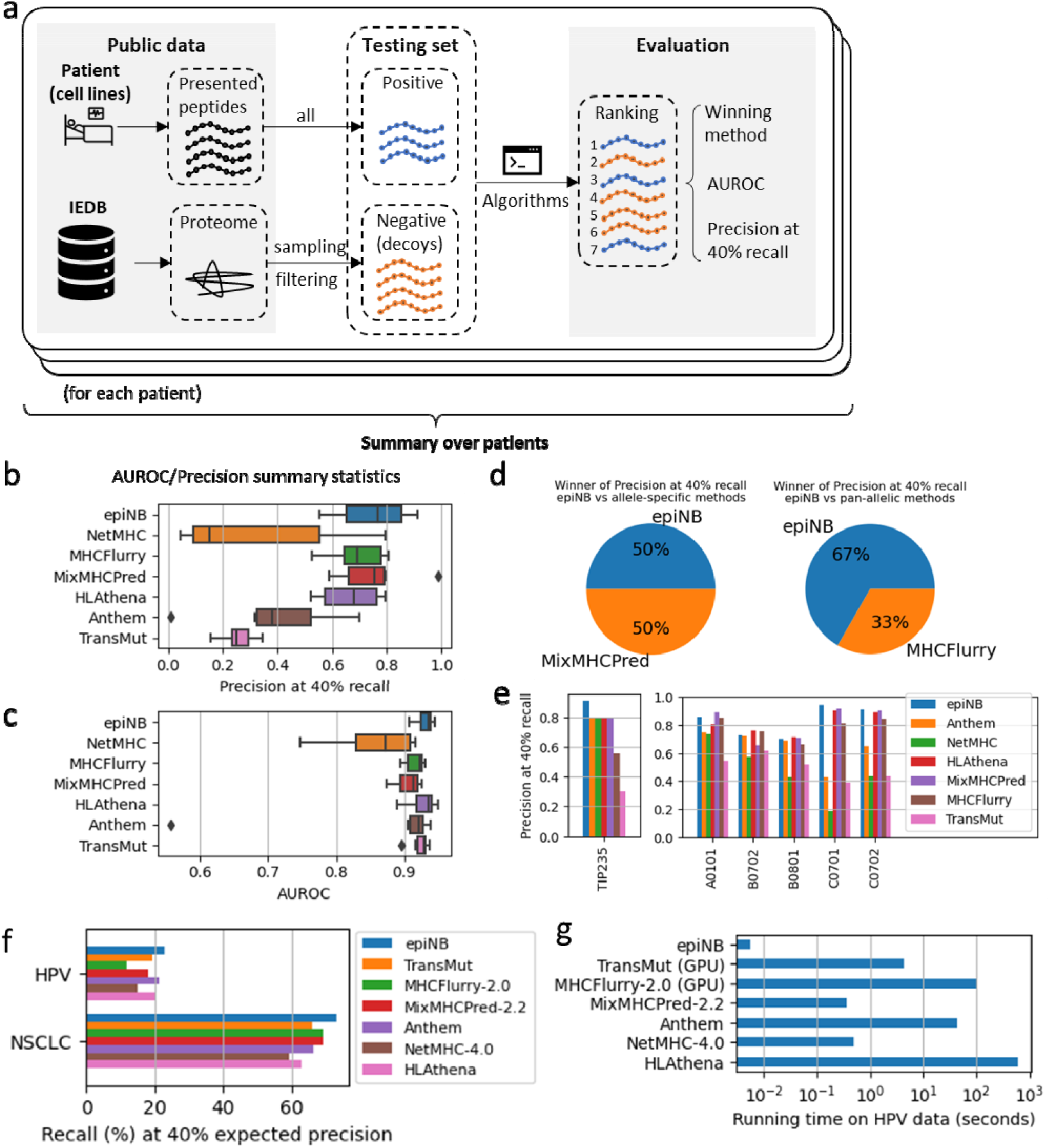
Benchmarking on patient datasets. (a) Workflow of benchmarking on real data (b-c) precision at 40% recall (b) and area under receiver operational curve (AUROC) (c). Higher is better. (d) The proportion of winners over all 95 alleles. More is better. (e) Performance of all methods on TIP235 (left) and its individual alleles (right, as benchmarked with IEDB). (f) Recall on HPV and NSCLC neoantigen data. (g) Running time of all methods on the HPV dataset (278 peptides).

The results (**Figure 4b,c**) show that epiNB has state-of-the-art performance in AUROC and precision. It performs better on a majority of patients than other allele-specific methods (**Figure 4d**) and shows comparable performance as pan-allelic methods. Further, we identified interesting examples such as TIP235, a pancreatic cancer xenograft model, with five distinct HLA alleles (HLA-A is homozygous). Although epiNB is not always the best performer on these alleles when benchmarked on IEDB, its performance on this dataset is outstanding (**Figure 4e**). Our further experiment (**Supplementary Figure 8b,c**) shows that the performance would be on par with other methods without the semi-supervised training step, which clearly illustrates the benefits of this strategy. These results demonstrate our semi-supervised method could address the real patient-generating data.

To assess the performance of methods on experimentally determined neoantigens, Chu et al. [15] curated two datasets including 232 experimentally verified non-small-cell lung cancer (NSCLC) neoantigens with corresponding HLA alleles and another 278 from HPV16 proteins E6 and E7. Because the dataset contains only positive examples, we show recall scores at the calibrated thresholds for 40% precision (**Supplementary Table 1** shows a list of thresholds and precisions calibrated on the IEDB datasets). Results on both datasets show a clear edge of epiNB in recognizing neoantigens (**Figure 4f**).

We measured the running time of all methods using the HPV dataset (278 peptides) on a machine equipped with a Ryzen 5 3600 CPU (6 cores@3.6-4.2GHz), a GTX 3080 GPU, and sufficient (48GB) memory for all methods to run without swapping. EpiNB, using only 6ms, is the fastest method, while HLAthena, the slowest, takes over 10 minutes. The speed would be more crucial for sweeping scans of possible mutations, easily scaling the problem up to hundreds of thousands of peptides. It is worth noting that the training process of epiNB is also inexpensive. It takes only 1.5 seconds to train all seven models for the alleles shown in the HPV dataset.

## Discussion

Personalized peptide prediction provides an important foundation for the development of immunotherapies for a number of human diseases, necessitating accurate and interpretable modeling of HLA-I-mediated antigen presentation. Advances in machine learning and computational power underlined over-parametrized models, introducing the trade-off of performance and interpretability. However, white-box models still play important roles, especially when training data is limited. Here, we illustrate that a Naïve Bayes based model tailored for MHC-I peptide presentation achieves top performance on patient data, neoantigens and oncovirus protein peptides, while keeping the interpretability of a white-box model. The method emphasizes a fast and easy training process without tweaking any hyperparameters, making it especially suitable for clinicians to train or finetune on patient data, which are usually behind the barrier of multiple convoluted privacy policies and laws.

The iMotifs identified by the model will help clinicians inspect the selected peptides, and researchers further understand the antigen presentation process. Biologically, the dependencies between peptide positions presented in the data can be caused by two processes--antigen presentation and evolution of the peptide sequence. As circumstantial evidence, our experiment with biology-based smoothing using the substitution probability from BLOSUM62 performed worse than simple Laplacian (additive/pseudo-counts) smoothing. The reason may be that BLOSUM62 matrix demonstrates the evolution aspect of the amino acid, but not antigen presentation. Thus, we conjecture that the dependency is an intrinsic property of antigen presentation, including pHLA binding. More structural evidence is needed to explain these dependencies.

Although pan-allelic methods enable extrapolation of well-characterized HLA alleles to the unknown ones and have the theoretical potential to perform better on the known ones by utilizing more training data, our comparison suggests that they generally do not perform better than allele-specific methods. The usage of data from monoallelic cell line also contributed to the high accuracy. Thus, to design a patient-specific cancer vaccine that is of high stake, it may still be beneficial to perform peptide elution and use an allele-specific model to perform the prediction. The pan-allelic methods, on the other hand, can be used in gaining insights into the binding process and interpreting the results.

The recent adaptation of Transformer as a pan-allelic prediction backend makes it possible to interpret the model by using the attention scores [15], which reveals the most important positions for binding, and the amino acids that tend to bind at those positions. However, the iMotifs identified and used by epiNB, presumably captured by the subsequent neural network layers, would be elusive to interpret. EpiNB provides a more definitive explanation for binding for clinicians’ reference.

The variance (or inter-quantile distance) in performances of each method over HLA alleles and patients is also worth noting. Detailed numbers suggest that each method has a clear edge in some cases that may not be purely random fluctuation. This may give an edge to ensemble models. Further studies may want to inspect these cases, which may help develop a more omnibus model.

In summary, we introduced a white-box model with state-of-the-art accuracy, emphasizing interpretability, and specifically having an edge on small samples. The tool can be used to extract insights for peptide binding and make predictions to facilitate personalized cancer vaccine design.

## Methods

### Feature selection based on mutual information

The full set of features *F* consists of three parts. The first two parts are shared among all HLA alleles. Firstly, there are nine single-position features, named 1, 2, 3, 4, and 5 for the first five positions, and 7, 8, 9, and 0 for the last four positions. This naming strategy does not change by the length of peptides. (Note that for 8-mers, position 5 and 7 will be the same.) We then calculate the mutual information of each pair of positions *l*(*f_i_*; *f_j_*) = *H*(*f_i_*) + *H*(*f_j_*) – *H*(*f_i_*, *f_j_*) on all training peptides from all alleles and keep the top ten combinations with the highest mutual information, namely pan-allelic features. For each allele, we also use the binders for it to recalculate the allele-specific mutual information, and also keep the top ten combinations, namely allele-specific features. If a feature is already in the pan-allelic features, the 11^th^, etc. will be added until ten distinct ones are selected. In total, there will always be 29 features.

### Feature selection based on Siamese Network

We implemented a feedforward network *S*. During training, the network *S* takes one sample *a* (anchor), a positive samples *p* from the same class (HLA allele or patient) as the anchor, and a negative sample from another class *n*, and find their embeddings *S*(*a*), *S*(*p*), *S*(*n*), respectively. It then minimizes the triplet loss max(∥*S*(*a*) – *S*(*p*)∥^2^ – ∥*S*(*u*) – *S*(*n*)∥^2^ + *a*, 0).

In the input layer, amino acids and their combinations are encoded as intergers 1~20 and 1~400 and further transformed into one-hot vector of length 20 or 400. The weight of the first layer is extracted to indicate the importance of the features.

### Peptide classification with positive-example-only learning

To allow for training without negative examples, we created a model based on Naïve Bayes formulation. By Bayesian law, the probability of a peptide being a binder is

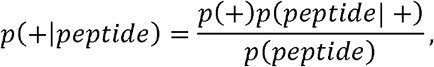

where *p*(+) is the prior probability of any peptide being a binder, *p(peptide*) is the probability of observing the given peptide, and *p*(*peptide*| +) is the probability of observing the given peptide in binders. The Naïve Bayes approach assumes that *p*(*peptide* | +) can be expressed as a product of a series of features Π_*f*∈*F*_ *p*(*f*| +), which are assumed to be mutually independent and bear equal weights. Under such an assumption, we have

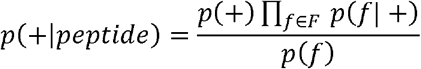

where *p*(*f*| +) can be estimated from training data. Here, we used the dataset generated in the HLAthena study [6], one of the cleanest datasets, which is generated from cell lines with single HLA alleles. Similarly,

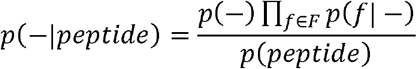

Because we do not have negative examples, the *p*(*feature*| −) are estimated from natural frequencies of AAs instead. *p*(+), *p*(−),and*p*(*peptide*) are hard to determine, but by noticing that *p*(+|*peptide*) + *p*(−|*peptide*) = 1, we can further deduce

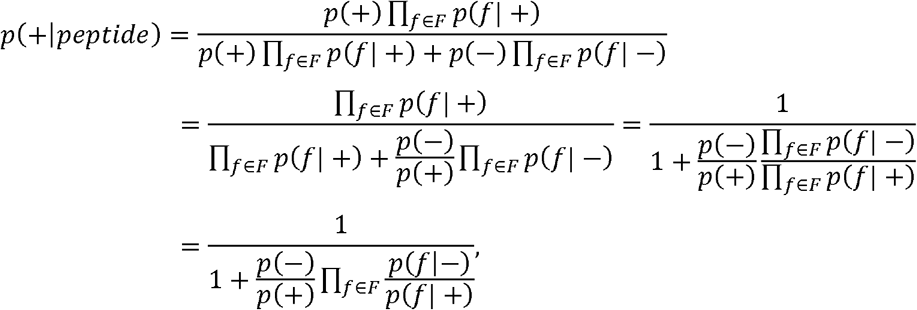

It is apparent that 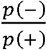 will only affect the value, but not the ordering of peptides, which is solely determined by the odds ratio 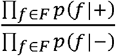. The actual calculation for 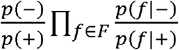 is performed on a logarithmic scale for better precision and speed. Due to numerical issues, it is usually helpful to set 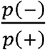 to a large number to avoid a large number of peptides receiving probability 1 and indistinguishable in the ordering step. In the experiments, 10^5^ – 1 was used (i.e., *p*(+) = 10^-5^).

In general, the odds ratio is less affected by numerical issues and is the recommended score to order the peptides.

### Semi-supervised personalized prediction

To deconvolve the mixture of peptides from multiple HLA alleles, we evaluate the probability of each peptide being a binder of each of the HLA allotypes of the patient and assign to it the top two predictions. These peptides are then added to the public data to form a larger training set, and retrain the models. When an HLA allotype of the patient does not have sufficient public data to train a model, we alternatively assign it to peptides that do not have a high probability for any other allotypes of the patient.

### Benchmarks

To construct the single-allelic testing dataset, we used all experimentally verified binders from IEDB as positive examples. We then randomly sample the human proteome in IEDB to create negative examples. For the dataset to be realistic, for each positive example, 99 negative examples are generated. Peptides that coincide with positive examples are removed. Substitutions are made to maintain the 1:99 ratio of positive and negative examples.

We also collected eluted peptides from 37 patient-derived xenograft or cell line samples. To test the performance of semi-supervised classification, we split the peptides from each patient into two halves as training as testing data in the abovementioned way, at a 1:99 ratio.

The non-small-cell lung cancer (NSCLC) neoantigen and HPV peptides dataset are directly from Chu et al. The NSCLC dataset includes 232 experimentally verified neoantigens. Each antigen is labeled with the corresponding HLA allele. The HPV dataset includes 278 from HPV16 proteins E6 and E7.

### Metrics

Two widely accepted metrics are used in the assessments. Precision at 40% recall is the precision 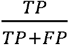 when the threshold is chosen to make the recall 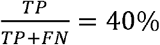. AUROC is the area under the receiver operating curve, the parametric curve of 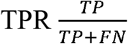 and 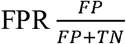.

On highly unbalanced datasets, precision at 40% recall is usually more indicative.

## Supplementary Materials

### Supplementary Note 1: Comparison of metrics

AUROC and precision at certain recall are both widely used metrics. However, the latter is more informative when the dataset is highly unbalanced, and the goal is to find a few positive examples. The definition of the three (as TPR is recall) are as follows.

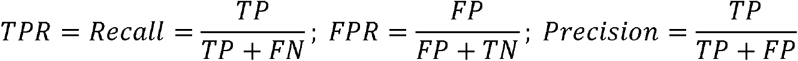

A relationship among these values can be deduced when the ratio of positive and negative examples are known.

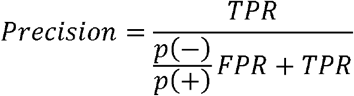

TPR (recall) and precision are more relevant to our task–select a few peptides that are likely to be binders. Here, recall indicates how many true binders we can recover, and precision tells how many decoys we have to include as a cost. FPR, on the other hand, tells the ratio of false positives and the number of decoys. It is not of immediate interest and can be misleading because a very small portion of many decoys can be disastrous. For example, when FPR=5% and TPR=90%, if *p*(–):*p*(+) = 100:1, the precision will only be 15%, not 100% - 5% = 95%. At this accuracy, a cancer vaccine would be technically impractical and financially prohibitive. This again indicates that a very large portion (FPR = 0.05~1.0) of the ROC has no practical value.

Using our result as an example, on HLA-A3001 (Supplementary Figure 3), epiNB achieves higher accuracy than HLAthena in almost all practically useful cases (precision > 0.2 and recall < 0.9). However, the AUROC of HLAthena (0.9830) is slightly higher than epiNB (0.9810). Since TPR is another name for recall, we can find the FPR at 0.9% FPR, which is clearly smaller than 0.1. This means that the small (and arguably useless for its low precision) portion (recall > 0.9) in the precision-recall curve actually decides > 90% of the AUROC.

In conclusion, we believe that precision at 40% recall is a more useful metric than AUROC for cancer vaccine design.

**Supplementary Table 1:** Thresholds for EpiNB at 99:1 Negative-Positive ratio

| Precision (median on all IEDB datasets) | Threshold for log odds |
| --- | --- |
| 90% | 29.0 |
| 80% | 23.0 |
| 70% | 19.0 |
| 60% | 15.0 |
| 50% | 11.5 |
| 40% | 7.5 |
| 30% | 3.0 |
| 20% | -3.0 |
| 10% | -12.0 |

\*The threshold may not be reliable for precision $> 50\%$ . Performance are significantly different on different alleles.

## Author contributions

GL, KC, HX, and SL conceptualized the project. SL, YC, and KC designed the algorithm. SL implemented the algorithm. GL, SL, and XJ collected data. SL benchmarked all methods. SL, XJ, KK, GL, and KC analyzed the data. All authors contributed to the manuscript preparation and approved the final version.

## Acknowledgments

This project has been made possible in part by grant RP180248 to KC from Cancer Prevention & Research Institute of Texas, grant U01CA247760 to KC and the Cancer Center Support Grant P30 CA016672 to PP from National Cancer Institute.

