## Supplementary Figures for "An interpretable ML model to characterize patient-specific HLA-I antigen presentation"

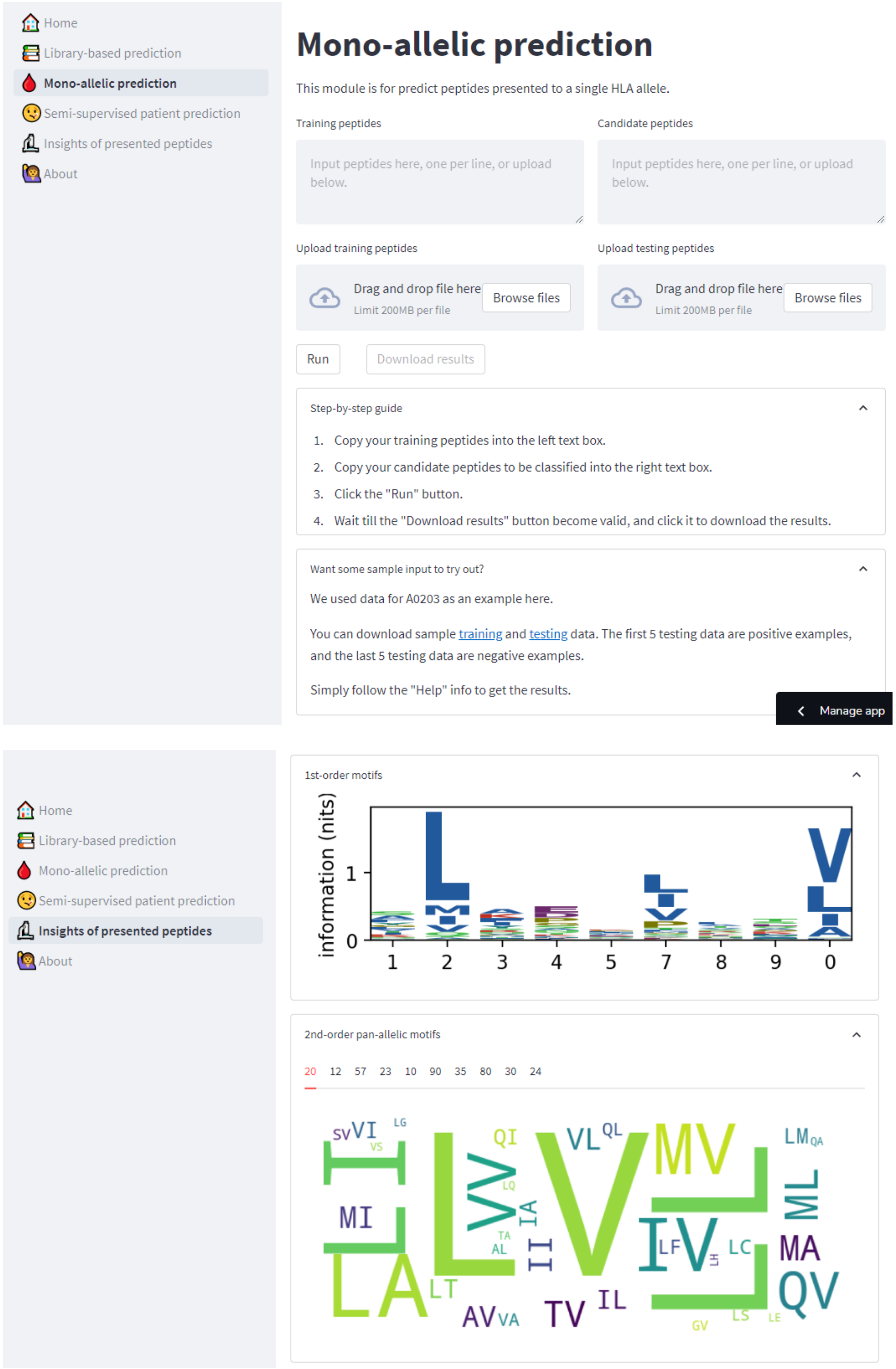


Supplementary Figure 1. EpiNB online portal for prediction and motif investigation

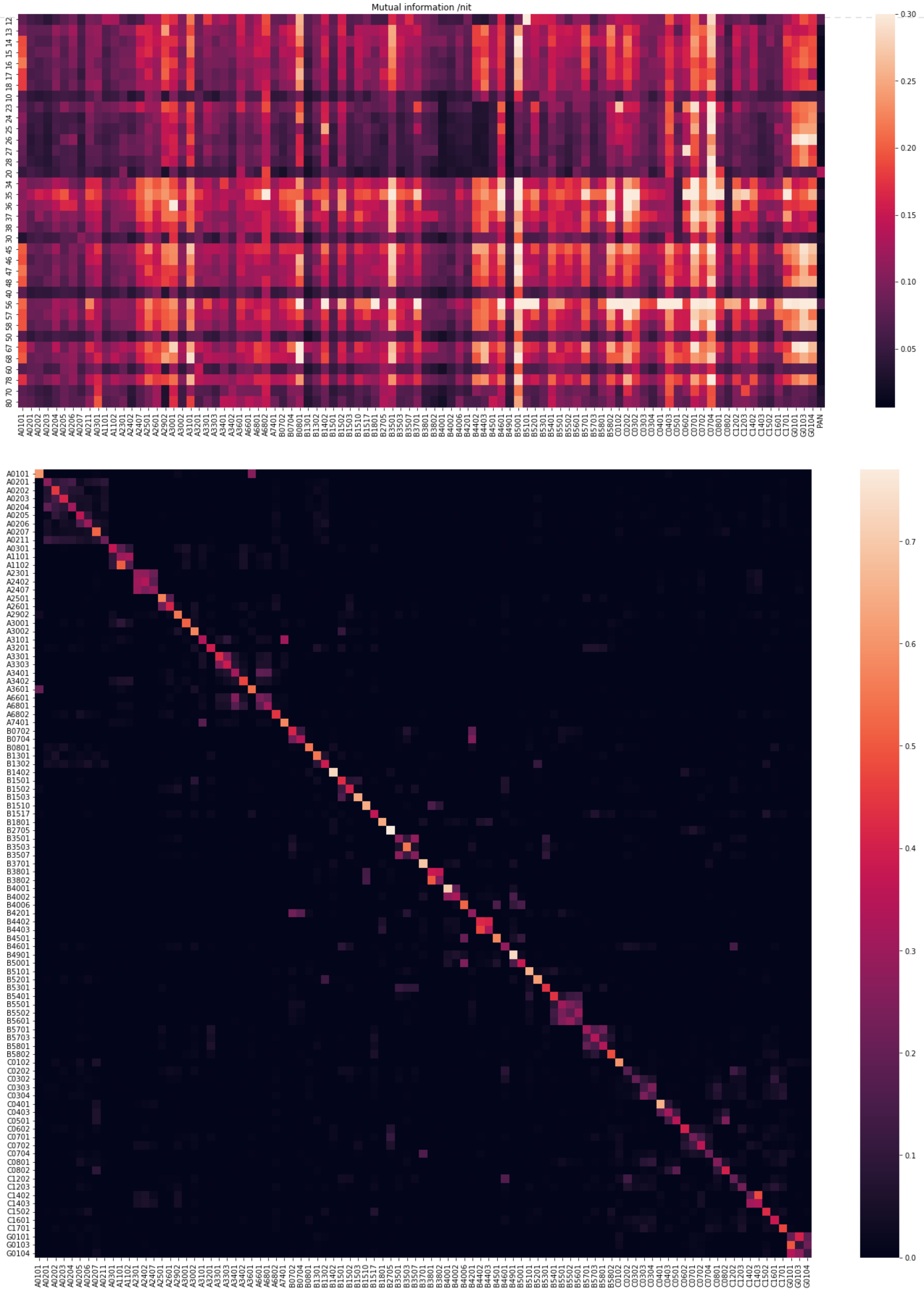


Supplementary Figure 2.

(a) Mutual information of combinations of position on each allele and on all alleles (last column named “PAN”).

(b) Confusion matrix of classification of allele origin of alleles. The monoallelic dataset was into two equal sets of training and testing data. EpiNB classifies the alleles of the peptides. We observed a decent performance of. Although some alleles appear to be harder to distinguish, they confirm the observation of HLA superfamilies in previous studies. The actual task on six alleles of patients would be significantly easier and a higher accuracy is expected.

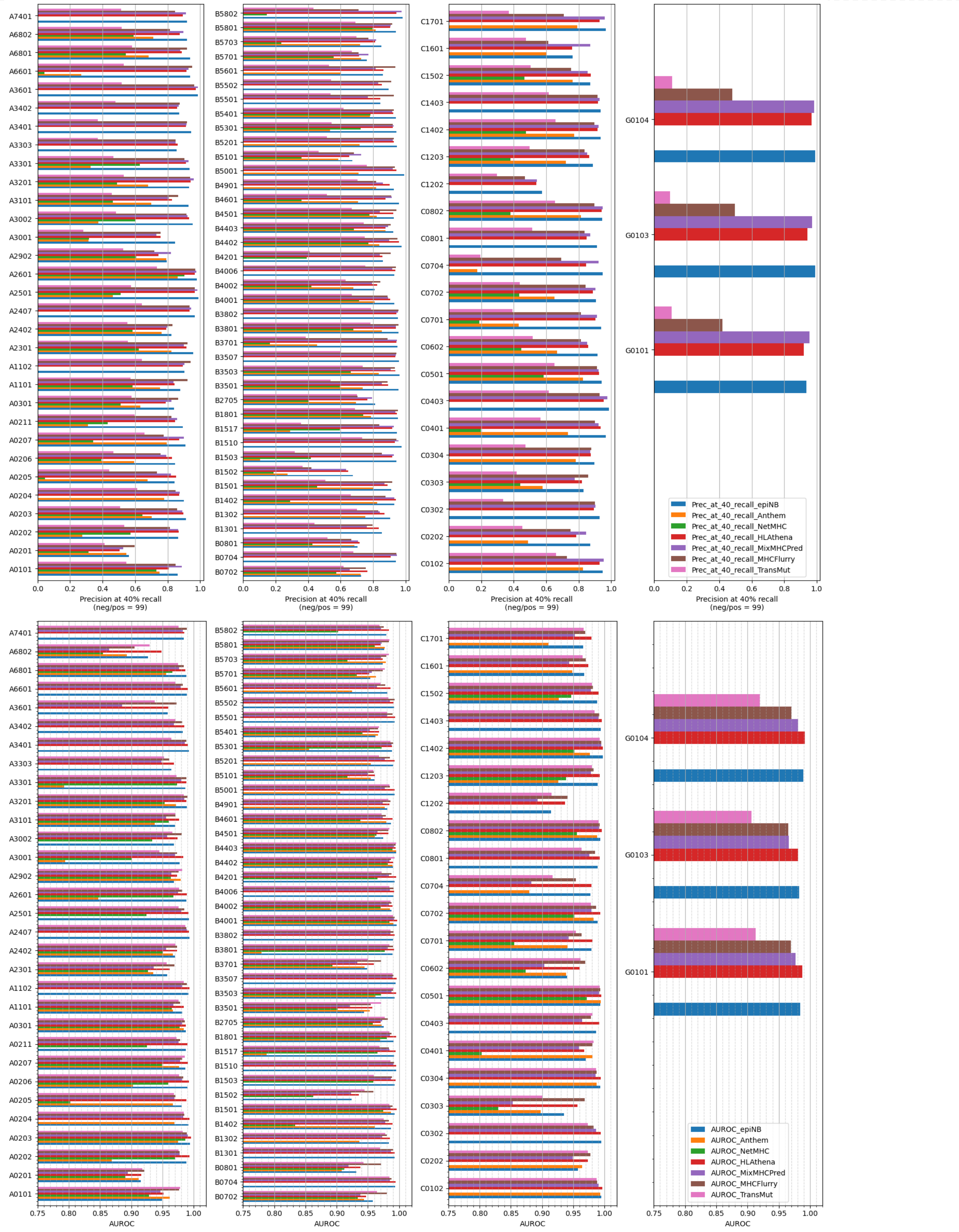


Supplementary Figure 3. Benchmarking on IEDB dataset

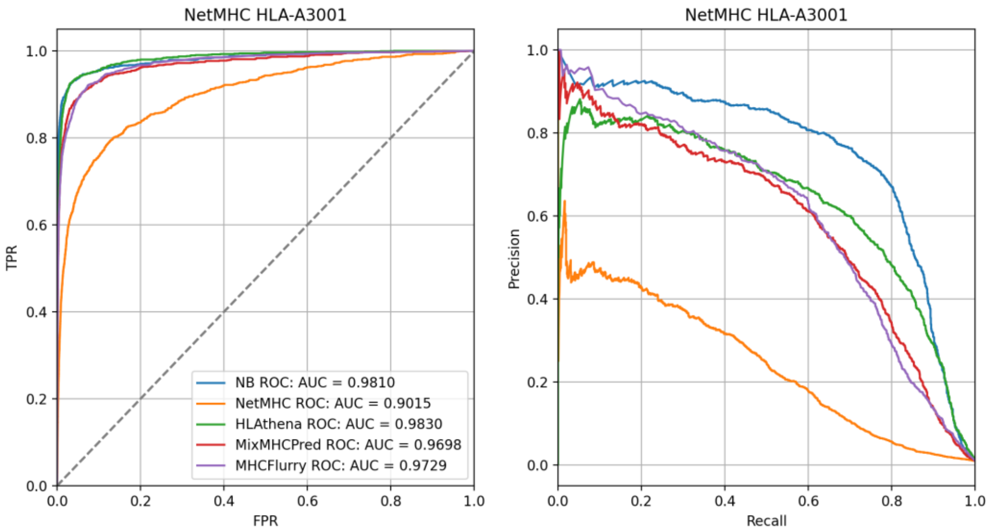


Supplementary Figure 4. Sample ROC and precision-recall curves from benchmarking on IEDB dataset a

| | A | C | D | E | F | G | H | I | K | L | M | N | P | Q | R | S | T | V | W | Y |
|---|---|---|---|---|---|---|---|---|---|---|---|---|---|---|---|---|---|---|---|---|
| **1** | -1.0 | -2.0 | 1.1 | 2.1 | -0.6 | -1.5 | 1.0 | -1.9 | -4.0 | -2.8 | -1.1 | -0.7 | -2.0 | -1.3 | -6.3 | -0.2 | -0.5 | -2.1 | -4.8 | 0.0 |
| **2** | -0.2 | -5.1 | -2.3 | -2.4 | -6.0 | -3.5 | -5.4 | 0.5 | -6.4 | -0.6 | -2.4 | -2.1 | -1.6 | -1.9 | -6.3 | -1.0 | 1.3 | 2.0 | -4.8 | -5.7 |
| **3** | 0.5 | -0.7 | -2.3 | -2.2 | 0.5 | -1.9 | -0.1 | 1.4 | 0.1 | -0.9 | 0.3 | -0.6 | 0.3 | -0.9 | -1.0 | -1.1 | -0.5 | 0.7 | -4.8 | 0.8 |
| **4** | -0.5 | -1.3 | 1.1 | 0.8 | -1.5 | 0.5 | 0.1 | -1.6 | 0.1 | -1.9 | -2.0 | -0.1 | 0.9 | 0.2 | -0.3 | -0.1 | 0.1 | -1.0 | -0.3 | -1.2 |
| **5** | -0.3 | -0.8 | -0.1 | 0.1 | 0.1 | 0.4 | 1.0 | -0.4 | 0.7 | -0.9 | -1.4 | -0.1 | 0.3 | 0.2 | 0.4 | 0.0 | -0.3 | -0.4 | -0.7 | -0.3 |
| **7** | -1.1 | -0.9 | -1.8 | -2.2 | 0.7 | -0.3 | -0.2 | 1.2 | -1.7 | 0.5 | 0.5 | -1.0 | -0.2 | -1.4 | -1.0 | -0.6 | 0.0 | 0.6 | -1.7 | 1.0 |
| **8** | 0.1 | -0.3 | -0.6 | 0.3 | -1.9 | -0.0 | 0.7 | -0.9 | -0.1 | -0.3 | -0.2 | 0.0 | -0.0 | 0.8 | 0.0 | 0.7 | 0.5 | -0.6 | -1.1 | -1.0 |
| **9** | 0.3 | 0.4 | -2.6 | 0.3 | -1.9 | -0.1 | 0.9 | -1.9 | 0.2 | -1.6 | -1.1 | -0.3 | -1.9 | 0.2 | -0.0 | 1.3 | 0.7 | -0.4 | -2.4 | -1.8 |
| **0** | -6.7 | -2.7 | -3.9 | -4.1 | 1.9 | -4.1 | -3.0 | -0.4 | -6.4 | 0.1 | -2.0 | -3.7 | -2.7 | -3.6 | -3.9 | -4.1 | -6.3 | -2.1 | 3.5 | 1.7 |

b

| | 20 | 12 | 57 | 23 | 10 | 90 | 35 | 80 | 30 | 24 |
|---|---|---|---|---|---|---|---|---|---|---|
| **0** | VW 5.4 | EV 4.0 | HY 2.1 | VI 3.4 | EW 5.3 | SW 4.7 | FH 2.4 | QW 4.3 | IW 4.6 | VD 3.2 |
| **1** | TW 5.0 | ET 3.4 | QI 2.0 | VV 2.8 | HW 4.8 | HW 4.5 | IH 2.3 | HW 4.2 | YW 4.2 | VE 3.0 |
| **2** | VF 3.8 | HV 3.1 | HF 2.0 | VY 2.7 | DW 4.7 | TW 4.2 | IK 2.3 | SW 4.1 | VW 4.1 | VP 2.8 |
| **3** | VY 3.8 | DV 3.0 | KY 2.0 | TI 2.6 | EF 3.9 | CW 3.9 | MH 2.2 | TW 4.0 | PW 4.0 | TD 2.4 |
| **4** | IW 3.8 | HT 2.7 | HM 2.0 | VA 2.5 | EY 3.9 | AW 3.8 | YG 2.0 | NW 3.8 | FW 3.9 | VG 2.4 |
| **...** | ... | ... | ... | ... | ... | ... | ... | ... | ... | ... |
| **395** | LE -4.2 | VL -4.2 | GA -4.0 | AS -4.0 | LG -4.2 | AA -4.1 | SS -3.9 | AA -4.1 | LE -4.2 | GA -4.0 |
| **396** | LV -4.2 | LG -4.2 | AS -4.0 | KL -4.1 | LS -4.2 | LE -4.2 | RL -4.0 | LE -4.2 | LV -4.2 | LI -4.1 |
| **397** | GL -4.2 | LS -4.2 | LK -4.1 | EL -4.2 | AL -4.4 | LV -4.2 | LD -4.0 | LG -4.2 | LG -4.2 | KL -4.1 |
| **398** | LG -4.2 | AL -4.4 | LE -4.2 | LE -4.2 | LA -4.4 | LG -4.2 | GA -4.0 | LS -4.2 | LS -4.2 | LV -4.2 |
| **399** | LA -4.4 | LL -4.6 | LA -4.4 | LG -4.2 | LL -4.6 | LA -4.4 | AL -4.4 | LA -4.4 | LA -4.4 | GL -4.2 |

c

| | 45 | 37 | 34 | 58 | 78 | 38 | 89 | 59 | 48 | 79 |
|---|---|---|---|---|---|---|---|---|---|---|
| **0** | DH 2.1 | II 2.8 | YD 2.4 | HQ 2.4 | IQ 2.0 | IQ 2.1 | QS 2.2 | HT 2.3 | DQ 2.1 | YC 2.5 |
| **1** | WH 2.0 | IF 2.4 | ID 2.3 | HH 2.3 | YQ 2.0 | IH 2.0 | HS 2.1 | CH 2.1 | DS 2.0 | VH 2.4 |
| **2** | EK 1.9 | FM 2.2 | IE 2.2 | RQ 1.6 | IC 1.8 | YH 1.9 | SS 2.0 | HC 2.1 | DH 1.9 | YS 2.4 |
| **3** | GP 1.9 | IY 2.1 | IQ 2.1 | HS 1.6 | YS 1.8 | YN 1.9 | EH 2.0 | HS 2.1 | HC 1.7 | IS 2.2 |
| **4** | DK 1.8 | FY 2.0 | AP 1.9 | NH 1.4 | IH 1.8 | FH 1.9 | HH 1.8 | KS 1.9 | PH 1.7 | FS 2.1 |
| **...** | ... | ... | ... | ... | ... | ... | ... | ... | ... | ... |
| **395** | LI -4.1 | LD -4.0 | LV -4.2 | AD -3.8 | KA -3.9 | LI -4.1 | LD -4.0 | LI -4.1 | LK -4.1 | LD -4.0 |
| **396** | IL -4.1 | EA -4.0 | VL -4.2 | LD -4.0 | EA -4.0 | LK -4.1 | DL -4.0 | IL -4.1 | LE -4.2 | AV -4.0 |
| **397** | LV -4.2 | GA -4.0 | GL -4.2 | VA -4.0 | SA -4.0 | EL -4.2 | LI -4.1 | LV -4.2 | LV -4.2 | LI -4.1 |
| **398** | LS -4.2 | SA -4.0 | SL -4.2 | IL -4.1 | KL -4.1 | GL -4.2 | IL -4.1 | GL -4.2 | LA -4.4 | EL -4.2 |
| **399** | LA -4.4 | LE -4.2 | LA -4.4 | LI -4.1 | EL -4.2 | LL -4.6 | VL -4.2 | LL -4.6 | LL -4.6 | LL -4.6 |

Supplementary Figure 5. An example of epiNB log odds table for A2301, including 9 positions (a), 10 pan-allelic features (b), and 10 allele-specific features (c). A sample result is shown in Fig. 2f.

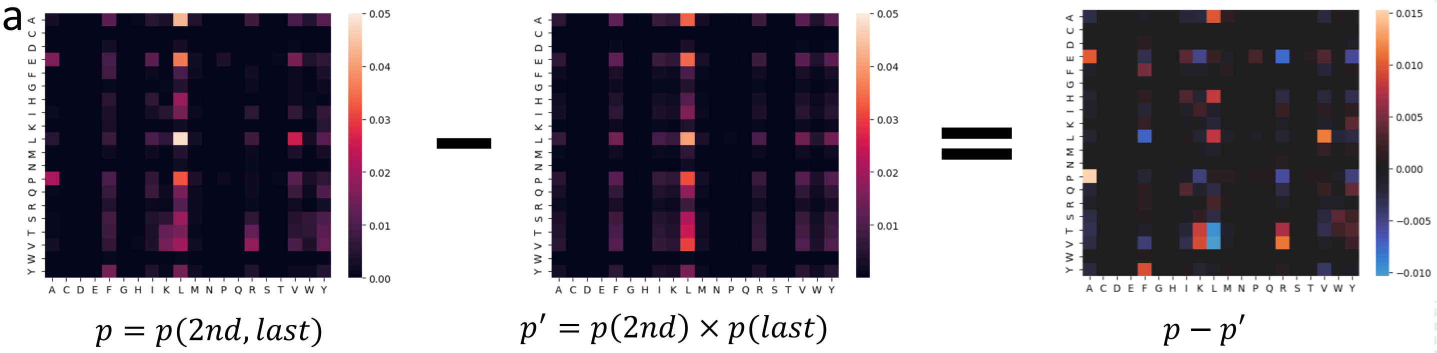


b

| | 20 | 12 | 57 | 23 | 10 | 90 | 35 | 80 | 30 | 24 |
|---|---|---|---|---|---|---|---|---|---|---|
| 0 | PA 1.5 | AE 0.9 | VV 0.9 | LP 0.5 | ER 1.7 | AA 0.5 | FG 0.6 | EL 0.4 | DL 1.2 | LE 0.8 |
| 1 | LV 1.2 | EV 0.9 | AA 0.8 | AD 0.5 | FL 0.8 | PK 0.4 | SV 0.5 | PL 0.3 | PL 0.7 | LD 0.6 |
| 2 | VR 1.1 | LP 0.7 | PP 0.7 | AA 0.4 | DR 0.8 | AL 0.3 | YG 0.4 | IK 0.3 | LK 0.2 | EL 0.4 |
| 3 | EA 1.0 | GE 0.6 | GG 0.7 | PR 0.4 | SK 0.4 | GR 0.3 | AP 0.3 | QL 0.3 | FK 0.2 | VD 0.3 |
| 4 | AL 1.0 | FA 0.5 | II 0.6 | PK 0.4 | YL 0.4 | EL 0.3 | TV 0.3 | VR 0.3 | SL 0.2 | SP 0.3 |
| … | … | … | … | … | … | … | … | … | … | … |
| 395 | PR -0.6 | EL -0.4 | VP -0.2 | PI -0.4 | AR -0.3 | LA -0.3 | AG -0.2 | LL -0.3 | LL -0.3 | ED -0.3 |
| 396 | ER -0.7 | KP -0.4 | IL -0.2 | EP -0.4 | FR -0.3 | AI -0.3 | YV -0.2 | ER -0.3 | DA -0.4 | AE -0.3 |
| 397 | LF -0.8 | EP -0.5 | PV -0.3 | LV -0.4 | EV -0.4 | PL -0.3 | IV -0.3 | PY -0.3 | YL -0.4 | EP -0.4 |
| 398 | TL -0.9 | FE -0.5 | VL -0.3 | ED -0.5 | DL -0.4 | LV -0.4 | FI -0.3 | VL -0.3 | DR -0.5 | LL -0.4 |
| 399 | VL -1.0 | RA -0.6 | PI -0.4 | PD -0.5 | EL -1.0 | GL -0.4 | FV -0.3 | IL -0.4 | FL -0.6 | PD -0.5 |

c

```
         -24        -14        -4         7 9        17        24  27         37        45 47         57     63  67    74  77
B*54:01  MRVTAPRTLL LLLWGALALT ETWA|GSHSMR YFYTAMSRPG RGEPRFIAVG YVDDTQFVRF DSDAASPRGE PRAPWVEQEG PEYWDRNTQI YKAQAQTDRE SLRNLRGYY
B*07:02  -L-M----V- ---SA----- ----|------ ----SV---- -------S-- ---------- --------E- -----I---- ---------- ---------- ---------
B*08:01  -L-M----V- ---SA----- ----|------ --D------- -------S-- ---------- --------E- -----I---- ---------- F-TNT----- ---------
B*14:02  -L-M----V- ---SA----- ----|------ -----V---- -------S-- ---------- --------E- -----I---- ---------- C-TNT----- ---------
B*55:01  ---------- ---------- ----|------ ---------- ---------- ---------- --------E- -----I---- ---------- ---------- ---------
B*55:02  ---------- ---------- ----|------ ---------- ---------- ---------- --------E- -----I---- ---------- ---------- ---------
B*56:01  ---------- ---------- ----|------ ---------- ---------- ---------- --------E- -----I---- ---------- ---------- ---------
           87       95 97         107    114 117   123   127        137        147        157        167        177        187
B*54:01  N QSEAGSHTWQ TMYGCDLGPD GRLLRGHNQL AYDGKDYIAL NEDLSSWTAA DTAAQITQRK WEAARVAEQL RAYLEGTCVE WLRRYLENGK ETLQRADPPK THVTHHPI
B*07:02  - --------L- S-----V--- -------D-Y ---------- ----R----- ---------- -----E---R ------E--- ---------- DK-E------ --------
B*08:01  - --------L- S-----V--- ---------Y ---------- ----R----- ---------- ---------D ---------- ---------- D--E------ --------
B*14:02  - --------L- W-----V--- ------Y--F ---------- ---------- ---------- -----E---- ---------- ----H----- ---------- --------
B*55:01  - ---------- ---------- ---------- ---------- ---------- ---------- -----E---- ---------- ---------- ---------- --------
B*55:02  - ---------- ---------- ---------- ---------- ---------- ---------- ---------- ---------- ---------- ---------- --------
B*56:01  - ---------- ---------- ---------- ---------- ---------- ---------- ---------- ------L--- ---------- ---------- --------
            197        207        217        227        237        247        257        267        277        287        297
B*54:01  SD HEATLRCWAL GFYPAEITLT WQRDGEDQTQ DTELVETRPA GDRTFQKWAA VVVPSGEEQR YTCHVQHEGL PKPLTLRWEP SSQSTIPIVG IVAGLAVLAV VVIGAVV
B*07:02  -- ---------- ---------- ---------- ---------- ---------- ---------- ---------- ---------- -----V---- ---------- -------
B*08:01  -- ---------- ---------- ---------- ---------- ---------- ---------- ---------- ---------- -----V---- ---------- -------
B*14:02  -- ---------- ---------- ---------- ---------- ---------- ---------- ---------- ---------- -----V---- ---------- -------
B*55:01  -- ---------- ---------- ---------- ---------- ---------- ---------- ---------- ---------- ---------- ---------- -------
B*55:02  -- ---------- ---------- ---------- ---------- ---------- ---------- ---------- ---------- ---------- ---------- -------
B*56:01  -- ---------- ---------- ---------- ---------- ---------- ---------- ---------- ---------- ---------- ---------- -------
             307        317        327        337
B*54:01  ATV MCRRKSSGGK GGSYSQAASS DSAQGSDVSL TA
B*07:02  -A- ---------- --------C- ---------- --
B*08:01  -A- ---------- --------C- ---------- --
B*14:02  -A- ---------- ---------- ---------- --
B*55:01  --- ---------- ---------- ---------- --
B*55:02  --- ---------- ---------- ---------- --
B*56:01  --- ---------- ---------- ---------- --
```

Supplementary Figure 6. EpiNB improves interpretability of peptide binding predictions
(a) Heatmaps showing the frequency of combinations of AAs, corresponding to Figure 3a-c.
(b) Table showing the highly surplus and deficient combinations of AAs.
(c) Sequence alignment for pocket B: 9,24,45,63,66,67 (yellow) and F: 74,77,95,97,114,116, 123 (cyan). B0702, B0801, and B1402 are provided for comparison.

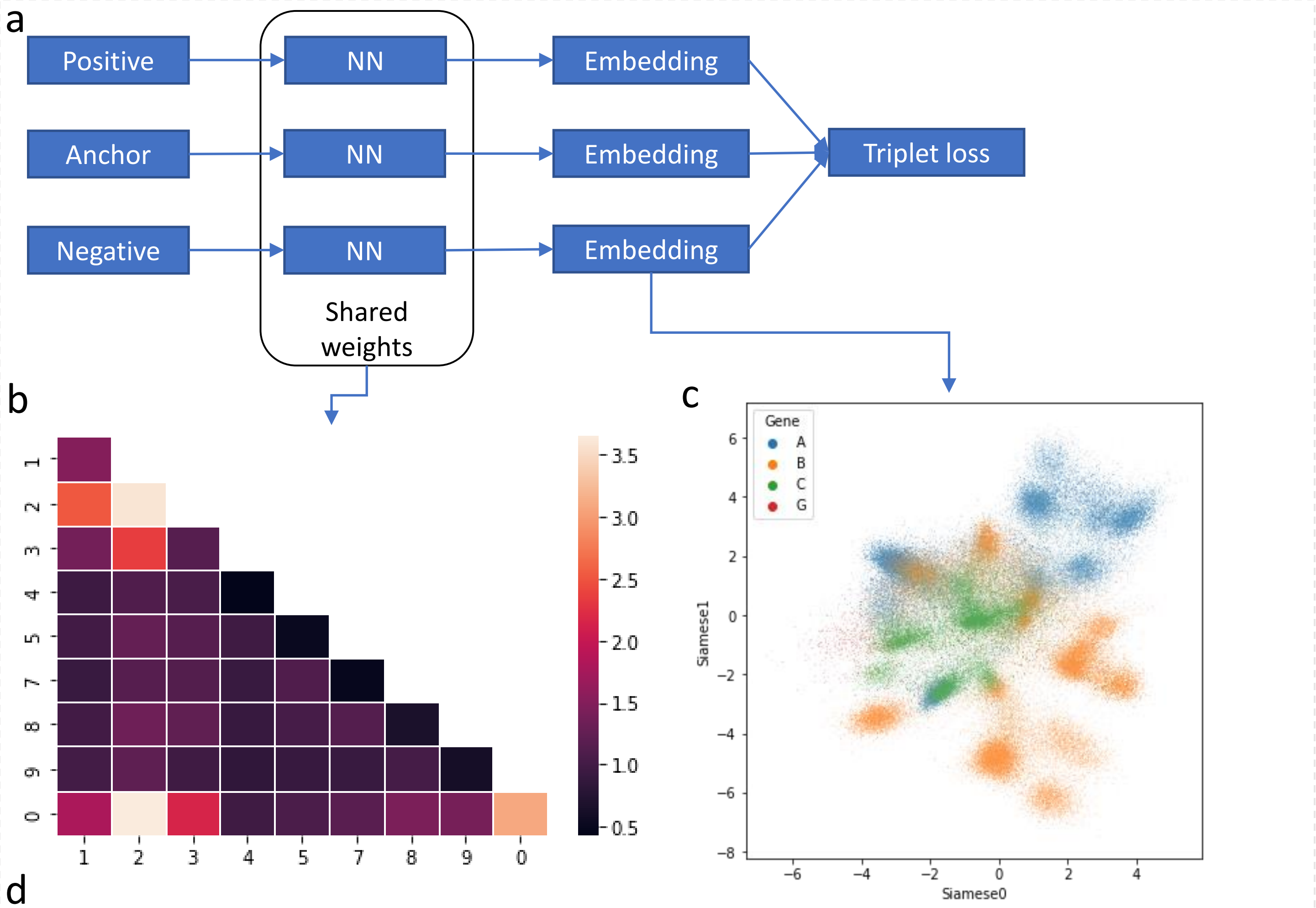


d

| Rank | Feature | Weight | Rank | Feature | Weight |
|---|---|---|---|---|---|
| 0 | **20** | 3.65 | 18 | **57** | 1.10 |
| 1 | **12** | 2.53 | 19 | 50 | 1.06 |
| 2 | **23** | 2.35 | 20 | 34 | 1.06 |
| 3 | **30** | 2.14 | 21 | 58 | 1.03 |
| 4 | **10** | 1.80 | 22 | 89 | 1.02 |
| 5 | **80** | 1.45 | 23 | 19 | 1.00 |
| 6 | **90** | 1.41 | 24 | 15 | 1.00 |
| 7 | 13 | 1.38 | 25 | 18 | 1.00 |
| 8 | 28 | 1.35 | 26 | 40 | 0.97 |
| 9 | 25 | 1.26 | 27 | 39 | 0.96 |
| 10 | 38 | 1.23 | 28 | 45 | 0.96 |
| 11 | 29 | 1.21 | 29 | 14 | 0.96 |
| 12 | 70 | 1.18 | 30 | 47 | 0.92 |
| 13 | **35** | 1.16 | 31 | 79 | 0.91 |
| 14 | 27 | 1.16 | 32 | 17 | 0.91 |
| 15 | 78 | 1.14 | 33 | 59 | 0.91 |
| 16 | 37 | 1.13 | 34 | 48 | 0.90 |
| 17 | **24** | 1.11 | 35 | 49 | 0.84 |

**Supplementary Figure 7**: Peptide interpretation and feature selection with Siamese network

(a) Network structure

(b) Weights of features. The diagonal represents the importance of single positions and the lower triangle represents the importance of combinatorial features.

(c) Output embedding of the network. Colors correspond to HLA-A/B/C/G. Different clusters are from different HLA alleles.

(d) Importance of combinatorial features.

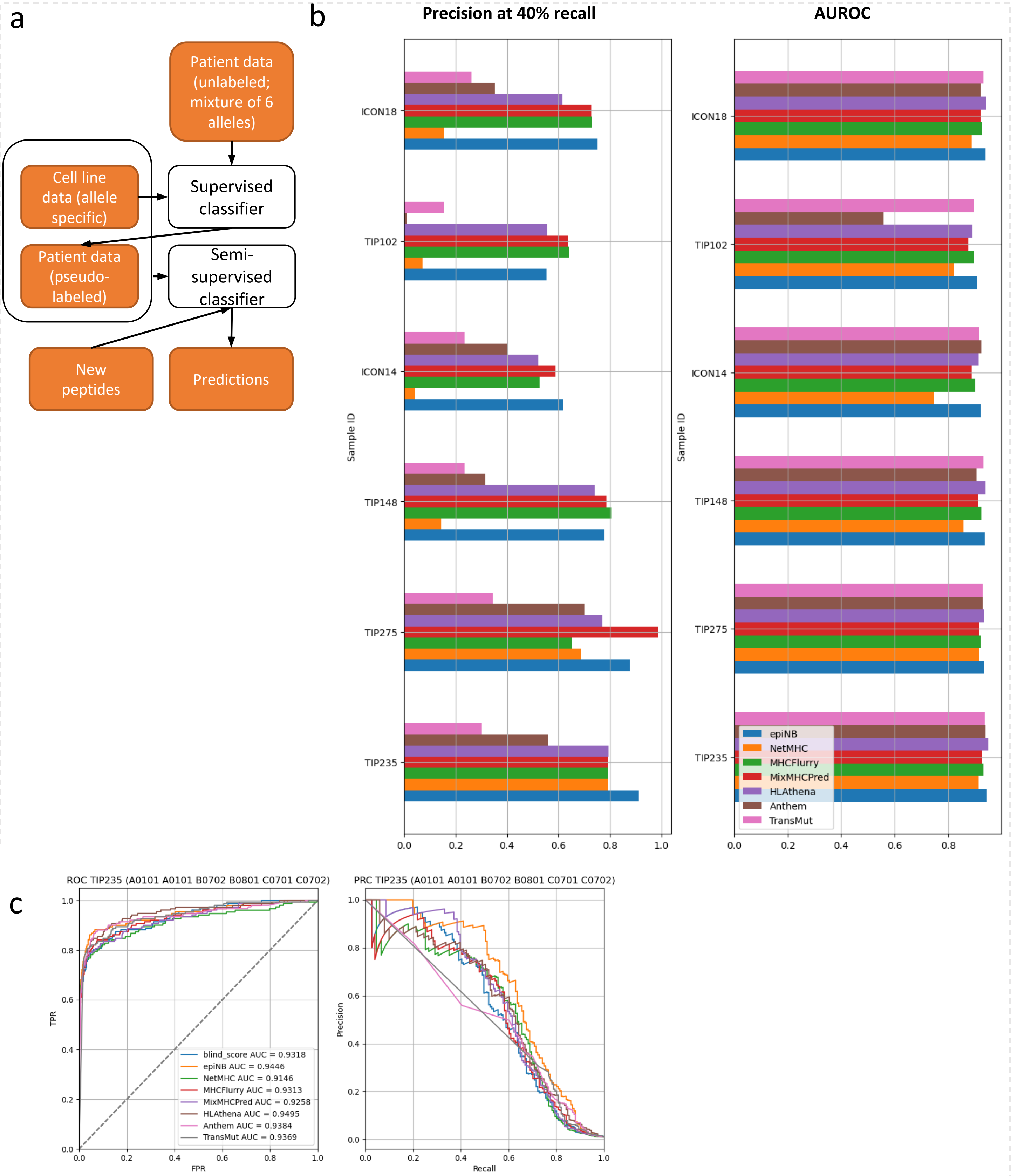


Supplementary Figure 8.

a. The pseudo-labeling approach for semi-supervised classification.

b. Precision at 40% recall and AUROC for all patient data.

c. Sample ROC and precision-recall curves for TIP235. Blind_score is from the epiNB trained without deconvolving the patient derived epitopes, and without the aid of public data.
